# Genetic variants associated with Alzheimer's disease confer different cerebral cortex cell-type population structure

**DOI:** 10.1101/266296

**Authors:** Zeran Li, Jorge L Del-Aguila, Umber Dube, John Budde, Rita Martinez, Kathleen Black, Qingli Xiao, Nigel J. Cairns, the Dominantly Inherited Alzheimer Network (DIAN), Joseph D. Dougherty, Jin-Moo Lee, John C Morris, Randall J. Bateman, Celeste M. Karch, Carlos Cruchaga, Oscar Harari

**Author notes:** To whom correspondence should be addressed: Carlos Cruchaga, PhD Associate Professor Department of Psychiatry The Hope Center Program on Protein Aggregation and Neurodegeneration Washington University, School of Medicine 425 S. Euclid Ave. BJC Institute of Heath. Box 8134 St. Louis, MO 63110, Oscar Harari, PhD Assistant Professor Department of Psychiatry Washington University School of Medicine 660 South Euclid Avenue B8134, St. Louis, MO 63110 //.

## Abstract

Alzheimer’s disease (AD) is characterized by neuronal loss and astrocytosis in the cerebral cortex. However, the effects of brain cellular composition are often ignored in high-throughput molecular studies. We developed and optimized a cell-type specific expression reference panel and employed digital deconvolution methods to determine brain cellular distribution in three independent transcriptomic studies. We found that neuronal and astrocyte proportions differ between healthy and diseased brains and also among AD cases that carry specific genetic risk variants. Brain carriers of pathogenic mutations in *APP*, *PSEN1* or *PSEN2* presented lower neurons and higher astrocytes proportions compared to sporadic AD. Similarly, the *APOE ε*4 allele also showed decreased neurons and increased astrocytes compared to AD non-carriers. On the contrary, carriers of variants in *TREM2* risk showed a lower degree of neuronal loss than matched AD cases in multiple independent studies. These findings suggest that genetic risk factors associated with AD etiology have a specific imprinting in the cellular composition of AD brains. Our digital deconvolution reference panel provides an enhanced understanding of the fundamental molecular mechanisms underlying neurodegeneration, enabling the analysis of large bulk RNA-seq studies for cell composition, and suggests that correcting for the cellular structure when performing transcriptomic analysis will lead to novel insights of AD.

## Introduction

Alzheimer’s Disease (AD) is a neurodegenerative disorder characterized clinically by gradual and progressive memory loss and pathologically by the presence of senile plaques (Aβ deposits) and neurofibrillary tangles (NFTs, Tau deposits) in the brain [41]. AD has a substantial but heterogeneous genetic component. Mutations in the amyloid-beta precursor protein (*APP*) and *Presenilin genes* (*PSEN1* and *PSEN2*) [21, 59] cause autosomal dominant AD (ADAD) which is typically associated with early-onset (<65 years). In contrast, the most common manifestation of AD presents late-onset (LOAD) and accounts for the majority of the cases (90-95%). Despite appearing sporadic in nature, a complex genetic architecture underlies LOAD risk. *APOE* ε4 is the most common genetic risk factor, increasing the risk in 3- to 8-fold [19]. In addition, recent whole genome and whole exome analysis have identified rare coding variants in *TREM2* [9, 32], *PLD3* [20], *ABCA7* [22, 63] and *SORL1* [26, 56] that are associated with AD and confer risk comparable to that of carrying one *APOE* ε4 allele. Besides age at onset, the clinical presentations of LOAD and ADAD are remarkably similar with an amnestic and cognitive impairment phenotype [57, 66]. A minor fraction of cases of ADAD have additional neurological findings, sometimes also seen in LOAD [57, 66].

Altered cellular composition is associated with AD progression and decline in cognition. Neuronal loss in the hippocampus is characteristic in the initial stages of AD, which could explain early memory disturbances [52, 71]. As the disease progresses, neuronal death is observed throughout the cerebral cortex. Furthermore, ∼25% of individuals who die by ∼75 years of age who were cognitively normal also presented substantial cerebral lesions that resemble AD pathology, including amyloid plaque, NFTs, and neuronal loss [37]. Thus, the identification of the brain cellular population structure is essential for understanding neurodegenerative disease progression [30]. However, stereology protocols for counting neurons can be tedious, require extensive training and are susceptible to technical artifacts which may lead to biased quantification of cell-type distributions [30].

Recently there has been a growing interest in understanding the transcriptomic changes attributed to AD [8, 16, 27, 46, 50, 53, 62, 72], as these may point to underlying molecular mechanisms of disease. These studies are typically designed to analyze the expression profiles of large cohorts ascertained from homogenized regions of the brain (e.g. bulk RNA-seq) of affected and control donors. However, bulk RNA-seq captures the gene expression of all of the constituent cells in the sampled tissue, and the altered cellular composition associated with AD has been reported to confound downstream analyses [62].

Digital deconvolution approaches enhance the interrogation of expression profiles to identify the cellular population structure of individual samples, alleviating the requirement of additional neurostereology procedures. These approaches have been developed, tested and applied to ascertain cellular composition altered in many traits [40, 51, 61, 75]. However, digital deconvolution has not been applied to identify the cellular population structure from RNA-seq from human brain tissue. Technical constraints restrict the dissociation of cells from the brains for very specific conditions [13, 73, 74]. Nevertheless, a limited number of RNA-seq from isolated cell populations from the brain have been generated [13, 73, 74]. Using these resources, we are now able to generate a reference panel for digital deconvolution of human brain bulk RNA-seq data.

We sought to investigate the cellular population structure in AD by analyzing RNA-seq from multiple brain regions of LOAD participants. To do so, we assembled a novel brain reference panel and evaluated the accuracy of digital deconvolution methods by analyzing additional cell-type specific RNA-seq samples and by creating synthetic admixtures with defined cellular distributions. Then we analyzed large cohorts of pathologically confirmed AD cases and controls (N = 613) and verified that it predicts cellular distribution patterns consistent with neurodegeneration. Finally, we generated RNA-seq from the parietal lobe of participants from the Knight-ADRC [39], including non-demented controls, LOAD cases, with enriched proportions of carriers of high-risk coding variants associated with AD, and also ADAD from the Dominantly Inherited Alzheimer Network [23] (DIAN). We compared the cell composition in ADAD and LOAD; and also evaluated differences among carriers of coding high-risk variants in *PLD3, TREM2* and *APOE* ε4. Our findings indicate that cell-type composition differs among carriers of specific genetic risk factors, which might be revealing distinct pathogenic mechanisms contributing to disease etiology.

## Materials and methods

### Subjects and Samples

#### DIAN and Knight-ADRC

Parietal lobe tissue of post-mortem brain was obtained with informed consent for research use and were approved by Washington University in St. Louis review board. RNA was extracted from frozen brain using Tissue Lyser LT and RNeasy Mini Kit (Qiagen, Hilden, Germany). RNA-seq Paired end reads with read length of 2×150bp were generated using Illumina HiSeq 4000 with a mean coverage of 80 million reads per sample (Table 1; **Table S1**). RNA-seq was generated for 19 brains from The Dominantly Inherited Alzheimer Network (DIAN), 84 brains with late-onset AD and 16 non-demented controls from The Charles F. and Joanne Knight Alzheimer’s Disease Research Center (Knight ADRC) [39]. The clinical status of participants was neuropathologically confirmed [47]. We identified three additional participants from the Knight ADRC study with PSEN1 (A79V, I143T, S170F) mutations. CDR scores were obtained during regular visits throughout the study prior to the subject’s decease [48]. A range of other pathological measurement were collected during autopsy including Braak staging, as previously described [11].

**Table 1.** Demographics and disease status of cohorts from four brain bank resources.

|  | **Mayo<sup>a</sup>** | **MSBB<sup>b</sup>** | **DIAN** | **Knight-ADRC** |
| --- | --- | --- | --- | --- |
| **Sample Size** | 191 | 300 | 19 | 103 |
| **Age** | 83 ± 7.77 | 83.3 ± 7.55 | 50.6 ± 7.06 | 85.1 ± 9.78 |
| **% Male** | 45.5 | 36 | 68.4 | 38.8 |
| **% *APOE* ε4+** | 33.2 | 31.7 | 14.3 | 45.6 |
| **Brain weight** | - | - | 1187.7 ± 184.5 | 1138.1 ± 142.5 |
| **AD<sup>c</sup>** | 82 | 135 | 19 | 87 |
| **PA<sup>d</sup>** | 29 | 0 | 0 | 0 |
| **Control** | 80 | 85 | 0 | 16 |
| **CDR<sup>e</sup> = 0** | - | 40 | 0 | 13 |
| **CDR = 0.5** | - | 40 | 0 | 9 |
| **CDR = 1** | - | 30 | 2 | 11 |
| **CDR = 2** | - | 44 | 4 | 14 |
| **CDR = 3** | - | 146 | 1 | 56 |

<sup>a</sup> Mayo stands for Mayo Clinic.
<sup>b</sup> MSBB stands for Mount Sinai Brain Bank.
<sup>c</sup> AD stands for Alzheimer's Disease.
<sup>d</sup> PA stands for pathological aging (amyloid plaques but no tau tangles).
<sup>e</sup> CDR stands for clinical dementia rating for available samples.

RNA was extracted from frozen brain tissues using Tissue Lyser LT and RNeasy Mini Kit (Qiagen, Hilden, Germany) following the manufacturer’s instruction. RIN (RNA integrity) and DV200 were measured with RNA 6000 Pico Assay using Bioanalyzer 2100 (Agilent Technologies). The RIN is determined by the software on the Bioanalyzer taking into account the entire electrophoretic trace of the RNA including the presence or absence of degradation products. The DV200 value is defined as the percentage of nucleotides greater than 200nt. RIN and DV200 for all the samples can be found on **Table S1**. The yield of each sample is determined by the Quant-iT RNA Assay (Life Technologies) on the Qubit Fluorometer (Fisher Scientific). The cDNA library was prepared with the TruSeq Stranded Total RNA Sample Prep with Ribo-Zero Gold kit (Illumina) and then sequenced by HiSeq 4000 (Illumina) using 2×150 paired end reads at McDonnell Genome Institute, Washington University in St. Louis with a mean of 58.14 ± 8.62 million reads. Number of reads and other QC metrics can be found in **Table S1**.

#### Mayo Clinic Brain Bank

Mayo Clinic Brain Bank RNA-seq was accessed from the AMP-AD portal (synapse ID = 5550404; accessed January 2017) (Table 1). Paired end reads of 2×101 base pairs were generated by Illumina HiSeq 2000 sequencers for an average of 134.9 million reads per sample. Neuropathology criteria, quality control procedures, RNA extraction and sequencing details are explained elsewhere [8].

RNA-seq based transcriptome data was generated from post-mortem brain tissue collected from cerebellum (189 samples) and temporal cortex (191 samples) of Caucasian subjects [2, 8]. RNA was extracted using Trizol^®^ reagent and cleaned with Qiagen RNeasy. RIN measurement was performed with Agilent Technologies 2100 Bioanalyzer. Samples with RIN greater than 5 were included. Library was prepared by Mayo Clinic Medical Genome Facility Gene Expression and Sequencing Cores with TruSeq RNA Sample Prep Kit (Illumina).

#### Mount Sinai Brain Bank

Mount Sinai Brain Bank RNAseq study was downloaded from the AMP-AD portal (synapse ID = 3157743; accessed January 2017) (Table 1). Single end reads of 100 nucleotides was generated by Illumina HiSeq 2500 System (Illumina, San Diego, CA) for an average of 38.7 million reads per sample [5].

This dataset contains 1030 samples collected from four post-mortem brain regions of 300 subjects: anterior prefrontal cortex (BA10), superior temporal gyrus (BA22), parahippocampal gyrus (BA36), and inferior frontal gyrus (BA44). RNAseq was generated using the TruSeq RNA Sample Preparation Kit v2 and Ribo-Zero rRNA removal kit (Illumina, San Diego, CA) [3].

#### iPSC-derived neurons

We have generated and characterized human iPSC made from human fibroblasts using non-integrating Sendai virus carrying OCT3/4, SOX2, KLF4, and cMYC [65, 68]. iPSCs were plated in a v-bottom plate in neural induction media (StemCell Technologies; 65,000 per well) to form highly uniform neural aggregates. After 5 days, neural aggregates were transferred onto PLO/laminin-coated tissue culture plates. Neural rosettes formed over 5-7 days. The resulting neural rosettes were then isolated by enzymatic selection (StemCell Technologies) and cultured as neural progenitor cells (NPCs). NPCs were then differentiated culturing in neural maturation medium (neurobasal medium supplemented with B27, GDNF, BDNF, cAMP).

#### TRAP-seq mice

All animal procedures were performed in accordance with the guidelines of Washington University’s Institutional Animal Care and Use Committee. The Rosa26^fsTRAP^ mice (Gt(ROSA)26Sor^tm1(CAG^-EGFP/Rpl10a,-birA)Wtp) [76] (The Jackson Laboratory) were crossed with PV^Cre^ mice (Pvalb^tm1(cre)Arbr^) [35] (The Jackson Laboratory) to produce PV-TRAP mice directing expression of EGFP-L10a ribosomal fusion protein in parvalbumin (PV) expressing cells. Purification of cell-type specific mRNA by translating ribosome affinity purification (TRAP) was described previously [34] with modifications. Briefly, PV-TRAP mouse brain was removed and quickly washed in ice-cold dissection buffer (1× HBSS, 2.5 mM HEPES-KOH (pH 7.3), 35 mM glucose, and 4 mM NaHCO_3_ in RNase-free water). Barrel cortex was rapidly dissected and flash-frozen in liquid nitrogen, and then stored at −80 °C until use. Affinity matrix was prepared with 150 µl of Streptavidin MyOne T1 Dynabeads, 60 µg of Biotinylated Protein L, and 25 µg of each of GFP antibodies 19C8 and 19F7. The tissue was homogenized on ice in 1 ml of tissue-lysis buffer (20 mM HEPES KOH (pH 7.4), 150 mM KCl, 10 mM MgCl_2_, EDTA-free protease inhibitors, 0.5 mM DTT, 100 µg/ml cycloheximide, and 10 µl/ml rRNasin and Superasin). Homogenates were centrifuged for 10 min at 2,000 × *g*, 4 °C, and 1/9 sample volume of 10% NP-40 and 300 mM DHPC were added to the supernatant at final concentration of 1% (vol/vol). After incubation on ice for 5 min, the lysate was centrifuged for 10 min at 20,000 × *g* to pellet insolubilized material. Then 200 µl of freshly resuspended affinity matrix was added to the supernatant and incubated at 4 °C for 16–18 hours with gentle end-over-end mixing in a tube rotator. After incubation, the beads were collected with a magnet and resuspended in 1000 µl of high-salt buffer (20 mM HEPES KOH (pH 7.3), 350 mM KCl, 10 mM MgCl_2_, 1% NP-40, 0.5 mM DTT and 100 µg/ml cycloheximide), and collected with magnet as above. After 4 times of washing with high-salt buffer, RNA was extracted using Absolutely RNA Nanoprep Kit (Agilent Technologies) following manufacturer’s instruction. RNA quantification was measured using Qubit RNA HS Assay Kit (Life Technologies) and the integrity was determined by Bioanalyzer 2100 using an RNA Pico chip (Agilent Technologies). The cDNA library was prepared with Clontech SMARTer and then sequenced by HiSeq3000. Single end reads of 50 base pairs were generated for an average of 29.2 million reads per sample (24 samples).

#### iPSC-derived microglia

The data was accessed from the AMP-AD portal (Synapse ID syn7203233). Myeloid progenitors expressing CD14/CX3CR1 were generated within 30 days of differentiation. iPSC-derived microglia were able to phagocytose and responded to ADP by producing intracellular Ca^2+^ transients, whereas macrophages lacked such response. The differentiation protocol was highly reproducible across several induced pluripotent stem cell (iPSC) lines.

### RNA-seq QC and Alignment

FastQC was applied to DIAN and Knight-ADRC RNAseq data to perform quality check on various aspects of sequencing quality [58]. The DIAN and Knight-ADRC dataset was aligned to human GRCh37 primary assembly using Star (ver 2.5.2b) [24]. We used the primary assembly and aligned reads to the assembled chromosomes, un-localized and unplaced scaffolds, and discarded alternative haploid sequences. Sequencing metrics, including coverage, distribution of reads in the genome [4], ribosomal and mitochondrial contents and alignment quality, were further obtained by applying Picard CollectRnaSeqMetrics (ver 2.8.2) to detect sample deviation. Additional QC metrics can be found in **Table S1**.

Aligned and sorted bam files were loaded into IGV [55] to perform visual inspection of target variants. Samples carrying unexpected variants or missing expected variants were labeled as potential swapped samples. In addition, variants were called from RNA-seq following BWA/GATK pipeline [44, 45]. The identity of the samples was later verified by performing IBD analysis against genomic typing from GWAS chipsets.

### Expression quantification

We applied Salmon transcript expression quantification (ver 0.7.2) [54] to infer the gene expression for all samples included in the reference panel and participants in the Mayo, MSBB, DIAN and Knight-ADRC. We quantified the coding transcripts of *Homo Sapiens* included in the GENCODE reference genome (GRCh37.75). Similarly, we quantified the expression of the mice samples included in the reference panel using the Mus Musculus reference genome (mm10).

### Reference Panel

We assembled a cell-type specific reference panel from publicly available RNA-seq datasets comprised of both immunopanning collected or iPSC derived neurons, astrocytes, oligodendrocytes, and microglial cells from human and murine samples. For immunopanning collected cells, antibodies for cell-type specific antigens were utilized to bind and immobilize their targeted cell types in order to immunoprecipitate and purify each cell type from the suspensions [73]. cDNA synthesis was accomplished using Ovation RNA-seq system V2 (Nugen 7102) and library prepared with Next Ultra RNA-seq library prep kit from Illumina (NEB E7530) and NEBNext^®^ multiplex oligos from Illumina (NEB E7335 E7500). TruSeq RNA Sample Prep Kit (Illumina) was used to prepare library for paired-end sequence on 100ng of total RNA extracted from each sample. Illumina HiSeq 2000 Sequencer was used to sequence all libraries [73].

Both human adult temporal cortex tissue, collected from patients receiving neurological surgeries, and mice cells were disassociated, sorted and sequenced as described elsewhere [74], and deposited in the Gene Expression Omnibus GSE73721 and GSE52564. We also accessed neural progenitor cells (day 17) and mature human neurons (day 57 and 100) from Broad iPSC deposited in the AMP-AD portal [6] and neural progenitor cells and iPSC-derived neurons from [12]. Broad iPSC derived neurons accessed from AMP-AD portal were generated using an embryoid body-based protocol to differentiate into forebrain neurons [1]. Wild-type cells used in the protocol were obtained from UConn StemCell Core. RNA was purified using PureLink RNA mini-kit (Life Technologies) and libraries were prepared by Broad Institute’s Genomics Platform using TruSeq protocol. Please refer to **Table S2** for additional information.

#### Gene markers

The reference panel was assembled with samples from four distinct cell types. A redundant set of well-known cell-type markers was selected from the literature [74] (**Table S3**). Principal component analysis was performed on the reference panel using R function *prcomp* (version 3.3.3) to verify that the expressions of these gene were clustering samples by their cell types (**Fig S1b; Fig S2a**).

#### Inference of the cellular population structure

We ascertained alternative computation deconvolution algorithms implemented in the CellMix package (ver 1.6). Based on accuracy and robustness evaluation results we compared and reported the following three algorithms that outperformed the others: Digital Sorting Algorithm (named “DSA”) [75], which employs linear modeling to infer cell distributions; the method population-specific expression analysis (PSEA, also named meanProfile in CellMix implementation) [40] that calculates estimated expression profiles relative to the average of the marker gene list for each cell type [40]; and a semi-supervised learning method that employs non-negative matrix factorization (ssNMF in CellMix implementation) [29]. We tested additional methods which provided considerably lower accuracy (least-squares fit [7], quadratic programing [31]) or no significant difference (support vector regression [51] or latent variable analysis [17]) to the methods presented.

We selected the samples that provide the most faithful transcriptomic profile for their respective cell types by following a leave-one-out cross validation approach. We trained iteratively deconvolution models using all but one of the samples that was tested. Only samples predicted with a composition higher than 80% were kept for the reference panel (**Table S2; Fig S2b**).

### Accuracy and Robustness Evaluation

#### Chimeric validation

To emulate heterogeneous tissue with known and controlled cellular composition, we generated chimeric libraries pooling reads (to a total of 400,000) contributed from cell-type specific human donor samples (See **Table S2**). This process was repeated 720 times, using alternative samples from the reference panel to model each cell type. The proportion of reads that the libraries of neurons, astrocytes, oligodendrocytes and microglia provided to the chimeric libraries varied in predefined ranges (**Fig S3**). As a result, each of the chimeric libraries contained reads that followed 32 different distributions (neuronal reads contributed between 2 to 36% of reads, astrocytes between 22 to 76%, oligodendrocytes between 6 to 62% and microglia between 1 to 5%). Refer to **Table S4** for detailed description of the 32 different distributions. We quantified the chimeric reads using Salmon (v0.7.2) [54], and employed the samples that did not contribute reads to the chimeric library as reference panel for the deconvolution methods.

Overall, we quantified the expression of 23,040 (720 × 32) chimeric libraries. We evaluated the accuracy using the root-mean-square error (RMSE, **Equation 1** to compare the digital deconvolution cellular proportion estimates (method ssNMF) versus the defined proportion of reads specific to each of the chimeric libraries:

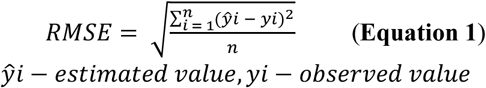

We also tested whether the deconvolution results were dominated by the expression of any specific gene, and ascertained the robustness of the transcriptomic signature we modeled for each cell type to any possibly altered gene expression. To do so, we performed the deconvolution analysis discarding one gene of the reference panel at a time and evaluated how these distributions differed in comparison to the full gene reference panel.

#### Statistical Analysis

We employed linear regression models to test the association between cell-type proportions and disease status (R Foundation for Statistical Computing, ver.3.3.3). We used stepwise discriminant analysis (stepAIC function of R package MASS, version 7.3-45) to determine significant covariates, and correct for confounding effects. We included RNA integrity number (RIN), batch, age at death and post-mortem interval (PMI) as covariates for the Mayo Clinic analyses. For Mount Sinai Brain Bank analyses, we corrected for RIN, PMI, race, batch and age at death. We also used linear-mixed models to perform multiple-region association analysis, employing random slopes and random intercepts grouping by observations and by donors [64], and correcting for the same covariates previously described.

To analyze the DIAN and Knight-ADRC studies we applied linear-mixed models (function lmer and Anova, R packages lme4 ver.1.1 and car ver.2.1, respectively), clustering at family level to ascertain the effect of the neuropathological status in the cell proportion, and corrected for RIN and PMI. For late-onset specific analyses we also corrected for age at death.

Cellular composition shown as proportions were plotted using R package ggplot2 (ver 2.2.1)

## RESULTS

### Study design

To infer cellular composition from RNA-seq, we firstly assembled a gene reference panel for neurons, astrocytes, oligodendrocytes and microglia. The panel was created by analyzing expression data from purified cell lines. We evaluated alternative digital deconvolution methods and selected the best performing for our primary analyses. We tested the digital deconvolution accuracy on induced pluripotent stem cell (iPSC) derived neurons/microglia cells and neuronal Translating Ribosome Affinity Purification followed by RNA-seq (TRAP-seq; Fig 1). Finally, we verified its accuracy by creating artificial admixture with pre-defined cellular proportions.

**Fig 1.**
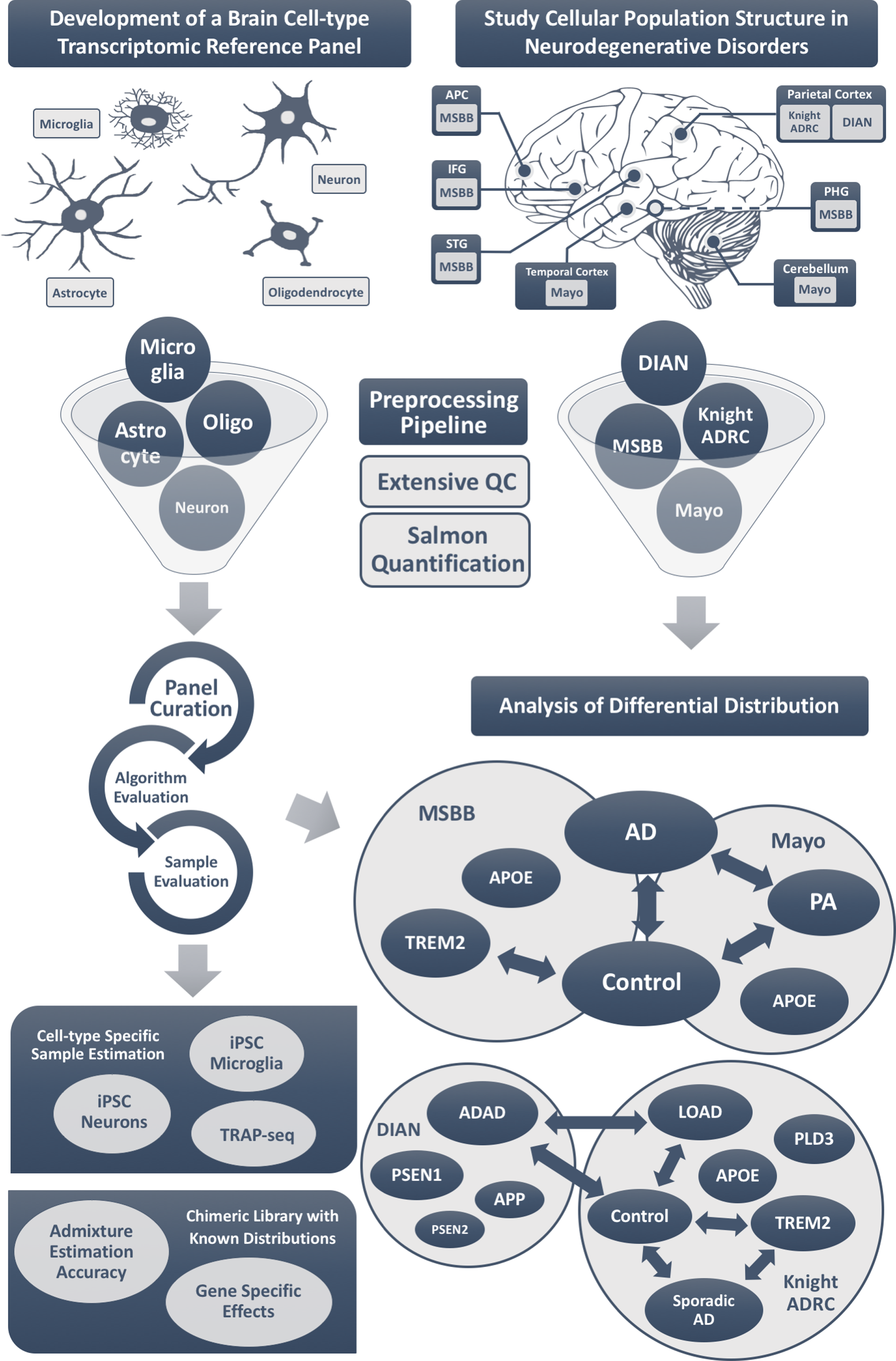
Study Design. Development of the brain cell-type transcriptomic reference panel (**left column**): the expression signatures of key cell types of the brain were curated by compiling publicly available RNA-seq data from neurons, astrocytes, oligodendrocytes and microglia. The panel was curated iteratively to retain only those samples that showed the most faithful expression signature, while evaluating alternative digital deconvolution methods. The accuracy of digital deconvolution to estimate brain cellular proportion was validated using additional cell-type specific samples, and also by generating chimeric libraries. To study cellular population structure in AD (**right column**), we accessed publicly available datasets from the Advanced Medicines Partnership-AD knowledge portal (AMP-AD), including Mayo Clinic and Mount Sinai Brain Bank datasets. In addition, we generated RNA-seq from participants of the Knight-ADRC and the Dominantly Inherited Alzheimer (DIAN) studies. These three studies generated RNA-seq data from pathological aging brains, Alzheimer Disease cases, and neuropath-free controls for a total of six cerebral cortex regions and cerebellum. We quantified the gene expression for all of the samples included in these studies using the same RNA-seq processing pipeline. Using digital deconvolution methods, we estimated the brain cellular proportions of the samples and compared the proportion between AD cases and controls. We study the cell structure of brains carriers of Mendelian pathological mutations and variants that confer high-risk to AD. Anterior prefrontal cortex – APC; superior temporal gyrus – STG; parahippocampal gyrus – PHG; inferior frontal gyrus – IFG; Mount Sinai Brain Bank – MSBB; Alzheimer Disease – AD; pathological aging – PA.

Once the deconvolution approach was optimized, we calculated the cell proportion in AD cases and controls from the different brain regions of Mayo and MSBB datasets. The RNA-seq data for the Mayo Clinic study (N = 191) [8] and Mount Sinai (MSSM) Brain Bank (MSBB; N = 300) [5] are deposited in the Advanced Medicines Partnership-AD (AMP-AD) knowledge portal (Synapse ID: syn5550404 and syn3157743; Table 1). The Mayo study includes RNA-seq from the temporal cortex and cerebellum for AD affected and non-demented controls, in addition to pathological aging participants (Fig 1). The MSBB also profiled four additional cerebral cortex areas: anterior prefrontal cortex - APC, superior temporal gyrus - STG, parahippocampal gyrus – PHG, and inferior frontal gyrus – IFG; Table 1; Fig 1). We restricted the case-control analysis to subjects with definite AD and autopsy confirmed controls. In addition, we generated RNA-seq from parietal lobe for participants of the Knight-ADRC (84 late-onset cases, carriers of genetic risk factors and 16 controls; **Table S1**) and the Dominantly Inherited Alzheimer Network (DIAN; 19 carriers of mutations in *APP*, *PSEN1*, *PSEN2*) (Table 1; Fig 1). We employed the same pipeline to process all of the samples in order to avoid any bias. Furthermore, RNA-seq from the Knight-ADRC and DIAN studies allowed us to compare the cell composition from ADAD vs LOAD brains, and similarly to test for differences in brain of controls, sporadic AD who do not carry any known high-risk variant, carriers of high-risk variants in *TREM2* (N = 20), *PLD3* (N = 33), and *APOE* e4 allele.

### Development of a reference panel to estimate brain cellular population structure

Due to limited availability of brain cell-type specific transcriptomic data, we compiled samples from different sources, including single-population RNA-seq from mice and human (immunopan-purified oligodendrocytes, neurons, astrocytes and microglia and iPSC-derived neurons and astrocytes) (**Table S2**).

We first tried to create a transcriptome wide reference panel by selecting the genes that are differentially expressed among cell types [17, 28, 51]. However, the species heterogeneity of the reference samples we compiled ruled out this attempt, as the principal component analyses (PCA) showed that differences between the human and mice donor samples dominated the transcriptome-wide effect (**Fig S1a**). For this reason, we curated a list of genes that have been described to tag these distinct cell types [14, 36, 74]. A visual inspection of the expression of these genes in the samples we compiled suggested a divergent transcriptomic profile among the cell types (**Fig S2a**). The PCA showed that their expression was sufficient to cluster samples of neurons, astrocytes, oligodendrocytes and microglia with their respective cell types, regardless of the species of the reference samples (**Fig S1b; Table S3**). We observed that some samples did not cluster with their expected cell types, and coincidently the leave-one-out cross-validation indicated that these samples had an expression signature that differed from the other samples of the same cell type. However, we found that all of these outliers correspond to samples not correctly purified or that were sequenced in early stages of differentiation (**Supplementary Results**). After discarding these samples, we assessed six digital deconvolution algorithms implemented in the CellMix package [28] and found that the semi-supervised non-negative matrix factorization [29] (ssNMF) calculated the most accurate estimates (see **Materials and methods).** Our final reference panel had a very high confidence to predict cell types with a mean predicted accuracy = 95.2%; s.d. = 4.3 (**Fig S2b**), and a root-mean-square error (RMSE) = 0.06 (**Table S5**).

### Optimization, validation and accuracy estimation of the reference panel and digital deconvolution method

Once we identified the optimal approach to perform digital deconvolution from brain RNA-seq, we benchmarked it by using three sets of independent pure cell populations and simulated chimeric libraries.

We firstly validated the accuracy to predict neuronal composition by generating RNA-seq for eight iPSC-derived cortical neurons (see **Materials and methods**). We observed an accurate prediction in these independent cell lines (mean neuronal proportion = 94.8% and s.d. = 1.1%; **Fig S4a**). We also ascertained the cellular composition of mRNA extracted from the barrel cortex neurons isolated by Translating Ribosome Affinity Purification (TRAP) in 24 mice. TRAP is a method that captures cell-type specific mRNA translation by purifying tagged ribosomal subunit and capturing the mRNA it bound to [34]. We observed an average of neuronal proportion = 96.7% and s.d. = 1.2% (**Fig S4b**). Similarly, we assessed the RNA-seq data generated for iPSC-derived microglia (N = 10) deposited in the AMP-AD portal (Synapse ID: -syn7203233) and inferred their cellular population structure, and observed a mean microglia proportion = 86.6% and s.d. = 7.1% (**Fig S4c**).

To evaluate the accuracy of digital deconvolution for measuring cell-type proportion from cell-type admixtures, we simulated RNA-seq libraries by pooling reads from individual cell types into well-defined proportions. We combined randomly sampled reads from neurons, astrocytes, oligodendrocytes and microglia to create chimeric libraries that mimic bulk RNA-seq from brain, but with a range of pre-defined cell-type distributions (**Fig S3**). We then quantified the gene expression for the chimeric libraries and inferred the cell-type distribution (employing for the reference panel samples that did not contribute reads to the chimeric libraries). This process was repeated 23,040 times, choosing distinct human samples to represent each cell type and varying the proportions in 32 alternative distributions (See methods and **Table S4**). The overall error (RMSE) compared to known proportions = 0.08.

Finally, we evaluated whether any gene included in the reference panel was dominating the inference of cell proportions. We re-calculated the cell-type distributions of the chimeric libraries, but dropping each of the genes from the reference panel one at a time. We observed a negligible difference between the cellular population structure inferred using the full reference and the gene-dropped panels (average RMSE = 0.022, s.d. < 0.01). In this way, we verified that the proportions inferred using the reference panel are not driven by the expression of a single gene. This reassured us the inference should be robust to any bias introduced by the potential association of a single gene included in the reference panel with a particular trait.

### Deconvolution of bulk RNA-seq of non-demented and AD brains shows a characteristic signature for neurodegeneration

Pathologically, AD is associated with neuronal death and gliosis specifically in the cerebral cortex. We evaluated whether we could exploit deconvolution methods using our reference panel to detect altered cellular population structure from the bulk RNA-seq, and whether this corresponded to known pathological alterations.

We initially analyzed the RNA-seq from the Mayo Clinic Brain Bank that includes bulk RNA-seq from the temporal cortex (TC) and cerebellum (CB) for 191 participants [8] (Table 1). In the TC, we observed a significant increase of astrocyte (β = 0.23; p = 5.01×10^−09^; Table 2; Fig 2; Table S6) in AD brains compared to controls brains. We also found a significant decrease of neurons (β = −0.17; p = 1.58×10^−07^; Table 2; Fig 2; Table S6) and oligodendrocytes (β = −0.07; p = 1.8×10^−02^; Table 2; **Fig S5; Table S6**). As expected, given the absence of pathology, we did not observe a significant change in the cell-type composition in the CB (Table 2).

**Fig 2.**
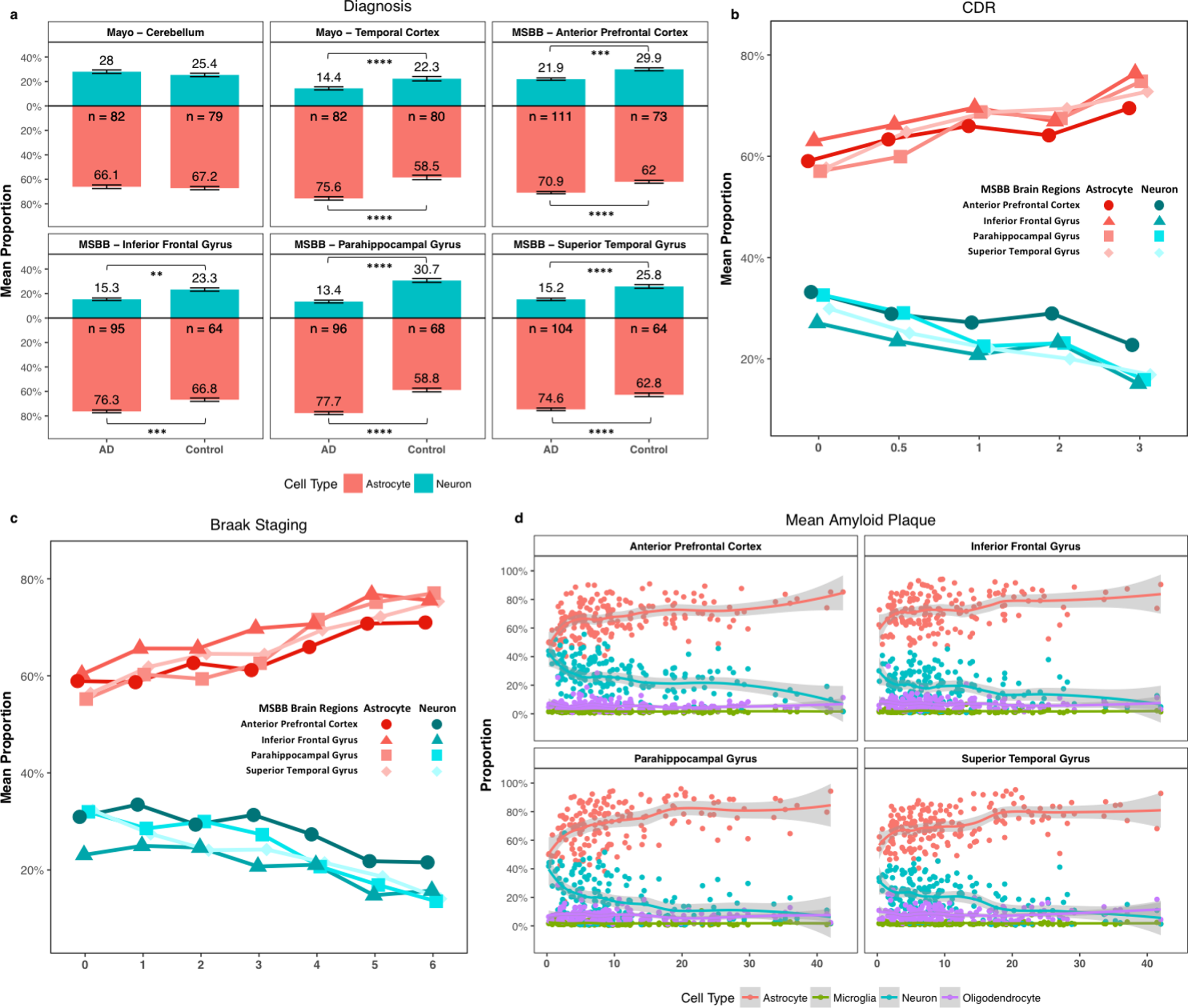
Cell-type distributions of the samples included in the Mayo Clinic and Mount Sinai Brain Bank. Mean neuronal (blue) and astrocytic proportion (red) for **a)** Alzheimer disease affected brains (AD) and controls (bars indicate standard deviations). The numbers of subjects for each group are shown below the x-axis. Distribution for additional clinical and pathological phenotypes reported for the Mount Sinai Brain Bank (MSBB): **b)** clinical dementia rating scores (CDR) and **c)** Braak and Braak staging. **d)** Brain cell-type proportions (x-axis) plotted against the mean number of amyloid plaque (values greater than 0; y-axis). Standard errors were depicted in shaded area with LOESS smooth curve fitted to cell-type proportions derived from deconvolution. (** P< 0.01; *** P< 1.0×10^−3^; and **** P< 1.0×10^−4^).

**Table 2.** Comparison of the cellular population structure (AD vs. neuropath-free controls) from the brains in the Mayo Clinic and Mount Sinai Brain Bank.

|  | Brain Regions | Sample Size | Neuron |  | Astrocyte |  | Oligodendrocyte |  | Microglia |  |
| --- | --- | --- | --- | --- | --- | --- | --- | --- | --- | --- |
|  | AD vs Control | N | Effect | P-value | Effect | P-value | Effect | P-value | Effect | P-value |
| Mayo | Cerebellum | 119 | -0.03 | $2.74 \times 10^{-01}$ | 0.05 | $8.65 \times 10^{-02}$ | -0.02 | $1.07 \times 10^{-01}$ | $-3.19 \times 10^{-04}$ | $9.19 \times 10^{-01}$ |
| | Temporal Cortex | 119 | -0.17 | $1.58 \times 10^{-07}$ | 0.23 | $5.01 \times 10^{-09}$ | -0.07 | $1.8 \times 10^{-02}$ | $-2.03 \times 10^{-03}$ | $5.48 \times 10^{-01}$ |
|  | **AD vs Control** |  |  |  |  |  |  |  |  |  |
| Mount Sinai Brain Bank | Anterior Prefrontal Cortex | 184 | -0.04 | $8.14 \times 10^{-04}$ | 0.06 | $8.11 \times 10^{-05}$ | -0.01 | $3.36 \times 10^{-02}$ | $-3.18 \times 10^{-03}$ | $1.12 \times 10^{-02}$ |
| | Superior Temporal Gyrus | 167 | -0.08 | $3.49 \times 10^{-07}$ | 0.1 | $1.45 \times 10^{-07}$ | -0.01 | $5.8 \times 10^{-02}$ | $-3.17 \times 10^{-03}$ | $5.78 \times 10^{-02}$ |
| | Parahippocampal Gyrus | 160 | -0.11 | $1.35 \times 10^{-08}$ | 0.13 | $5.48 \times 10^{-10}$ | -0.02 | $1.79 \times 10^{-03}$ | $-3.18 \times 10^{-03}$ | $1.35 \times 10^{-01}$ |
| | Inferior Frontal Gyrus | 159 | -0.04 | $3.12 \times 10^{-03}$ | 0.06 | $3.58 \times 10^{-04}$ | -0.01 | $4.39 \times 10^{-02}$ | $-3.98 \times 10^{-03}$ | $1.64 \times 10^{-02}$ |
|  | **Clinical Dementia Rating** |  |  |  |  |  |  |  |  |  |
| | Anterior Prefrontal Cortex | 184 | -0.02 | $9.38 \times 10^{-04}$ | 0.02 | $2.07 \times 10^{-04}$ | $-3.43 \times 10^{-03}$ | $1.25 \times 10^{-01}$ | $-1.46 \times 10^{-03}$ | $4.95 \times 10^{-03}$ |
| | Superior Temporal Gyrus | 167 | -0.03 | $1.87 \times 10^{-06}$ | 0.04 | $3.33 \times 10^{-07}$ | -0.01 | $2.1 \times 10^{-02}$ | $-1.02 \times 10^{-03}$ | $1.49 \times 10^{-01}$ |
| | Parahippocampal Gyrus | 160 | -0.04 | $8.56 \times 10^{-06}$ | 0.04 | $2.85 \times 10^{-06}$ | -0.01 | $8.7 \times 10^{-02}$ | $-1.94 \times 10^{-03}$ | $2.53 \times 10^{-02}$ |
| | Inferior Frontal Gyrus | 159 | -0.02 | $8.29 \times 10^{-05}$ | 0.03 | $1.4 \times 10^{-05}$ | $-4.64 \times 10^{-03}$ | $6.7 \times 10^{-02}$ | $-1.46 \times 10^{-03}$ | $3.11 \times 10^{-02}$ |
|  | **Braak Staging** |  |  |  |  |  |  |  |  |  |
| | Anterior Prefrontal Cortex | 173 | -0.01 | $1.21 \times 10^{-02}$ | 0.01 | $1.27 \times 10^{-03}$ | $-3.09 \times 10^{-03}$ | $2.77 \times 10^{-02}$ | $-7.04 \times 10^{-04}$ | $3.12 \times 10^{-02}$ |
| | Superior Temporal Gyrus | 158 | -0.02 | $2.22 \times 10^{-07}$ | 0.02 | $2.77 \times 10^{-07}$ | $-2.91 \times 10^{-03}$ | $1.17 \times 10^{-01}$ | $-5.47 \times 10^{-04}$ | $1.97 \times 10^{-01}$ |
| | Parahippocampal Gyrus | 147 | -0.02 | $1.83 \times 10^{-06}$ | 0.03 | $9.6 \times 10^{-08}$ | -0.01 | $1.49 \times 10^{-03}$ | $-3.71 \times 10^{-04}$ | $4.97 \times 10^{-01}$ |
| | Inferior Frontal Gyrus | 152 | -0.01 | $1.01 \times 10^{-02}$ | 0.01 | $8.56 \times 10^{-04}$ | $-3.55 \times 10^{-03}$ | $2.37 \times 10^{-02}$ | $-1.01 \times 10^{-03}$ | $1.74 \times 10^{-02}$ |
|  | **Mean Amyloid Plaques** |  |  |  |  |  |  |  |  |  |
| | Anterior Prefrontal Cortex | 184 | $-1.88 \times 10^{-03}$ | $3.6 \times 10^{-03}$ | $2.82 \times 10^{-03}$ | $1.03 \times 10^{-04}$ | $-7.99 \times 10^{-04}$ | $2.13 \times 10^{-03}$ | $-1.46 \times 10^{-04}$ | $1.72 \times 10^{-02}$ |
| | Superior Temporal Gyrus | 167 | $-4.2 \times 10^{-03}$ | $7.73 \times 10^{-08}$ | 0.01 | $4.63 \times 10^{-08}$ | $-6.08 \times 10^{-04}$ | $9.01 \times 10^{-02}$ | $-2.04 \times 10^{-04}$ | $1.5 \times 10^{-02}$ |
| | Parahippocampal Gyrus | 160 | $-4.96 \times 10^{-03}$ | $5.05 \times 10^{-09}$ | 0.01 | $1.26 \times 10^{-10}$ | $-9.99 \times 10^{-04}$ | $1.85 \times 10^{-03}$ | $-2.1 \times 10^{-04}$ | $2.58 \times 10^{-02}$ |
| | Inferior Frontal Gyrus | 159 | $-2.58 \times 10^{-03}$ | $3.82 \times 10^{-04}$ | $3.53 \times 10^{-03}$ | $1.96 \times 10^{-05}$ | $-7.41 \times 10^{-04}$ | $1.51 \times 10^{-02}$ | $-2.04 \times 10^{-04}$ | $1.26 \times 10^{-02}$ |

The cell-type proportions from AD cases and control were inferred from bulk RNA-seq using the ssNMF method. Effects of AD and associations with additional clinical and pathological phenotypes in cell-type distributions were estimated using linear regression model.

The distribution of microglia was similar in the TC and CB from AD and control brains (Table 2; **Fig S5**). The proportion of microglia was lower than any other cell types. The Mayo dataset also includes brains from individuals with pathological aging (PA; Table 1); which is neuropathologically defined by amyloid-beta (Aβ) senile plaque deposits but little or no neurofibrillary tau pathology [8, 49]. We observed a significant decrease of microglia proportion of PA brains compared to AD in both TC and CB (**Table S7; Fig S6**) [43]. Therefore, we speculated that the lack of changes in the AD microglial population was neither due to low statistical power nor the inability of our method to estimate the microglial proportions, but reflected unaltered neuropathological observations in AD brains.

We also analyzed data from the MSBB, which contains bulk RNA-seq for four additional cerebral cortex areas (APC, STG, PHG, IFG). Replicating our findings from the Mayo dataset we observed a significant decrease in neurons and increase in astrocytes in all four areas (Table 2; Fig 2; and Table S6). The strongest effect size was detected in the parahippocampal gyrus and superior temporal gyrus (p < 3.49×10^−07^) (Table 2; **Table S8**). Neuropathological studies have described that the parahippocampal gyrus in one of the first brain areas in which AD pathology occurs [10, 25, 69]. We also observed a significant and strong correlation between neuronal and astrocyte proportions and last ascertained clinical status (Clinical Dementia Rating - CDR), and number of amyloid plaques and Braak staging (Table 2; Fig 2; Fig S7).

### The cellular population structure differs between ADAD vs LOAD

While the loss of neurons is a common feature of AD, it is not clear whether the mechanism holds true across different forms of AD or AD cases carrying different genetic risk variants. Therefore, we investigated whether AD with distinct etiologies showed different cellular compositions. We generated RNA-seq data from the parietal lobe of participants enrolled in Knight-ADRC (84 LOAD, 3 ADAD, and 16 neuropath-free controls) and DIAN (19 ADAD) studies (Table 1; **Table S1**). We selected the LOAD and ADAD participants to match for CDR at death, brain weight and sex distributions (See **Table S1**).

Using digital deconvolution, we determined the cellular composition for these brains. We observed a significant decrease in neurons (β = −0.02, p = 2.66×10^−02^) and significant increase in astrocytes in AD (β = 0.03, p = 5.48×10^−03^) for the combined LOAD and ADAD brains compared to controls (Table 3; Fig 3; Table S9), consistent with our findings in the Mayo and MSBB datasets. Similarly, the joint analysis of the brains from Knight-ADRC and DIAN showed a significant association between the neuronal and astrocyte proportions and neuropathological measures (Braak staging: β = −0.03, p = 8.51×10-^06^ for neurons and β = 0.03, p = 3.83×10^−06^ for astrocytes; Table 3; Fig 3b) as well as for clinical measures (CDR: β = −0.02, p = 2.66×10^−02^ for neurons and β = 0.03 and p = 5.48×10^−03^ for astrocytes; Table 3; Fig 3c). We did not observe a significant difference in the compositions of microglia or oligodendrocytes (Table 3; **Fig S8**).

**Fig 3.**
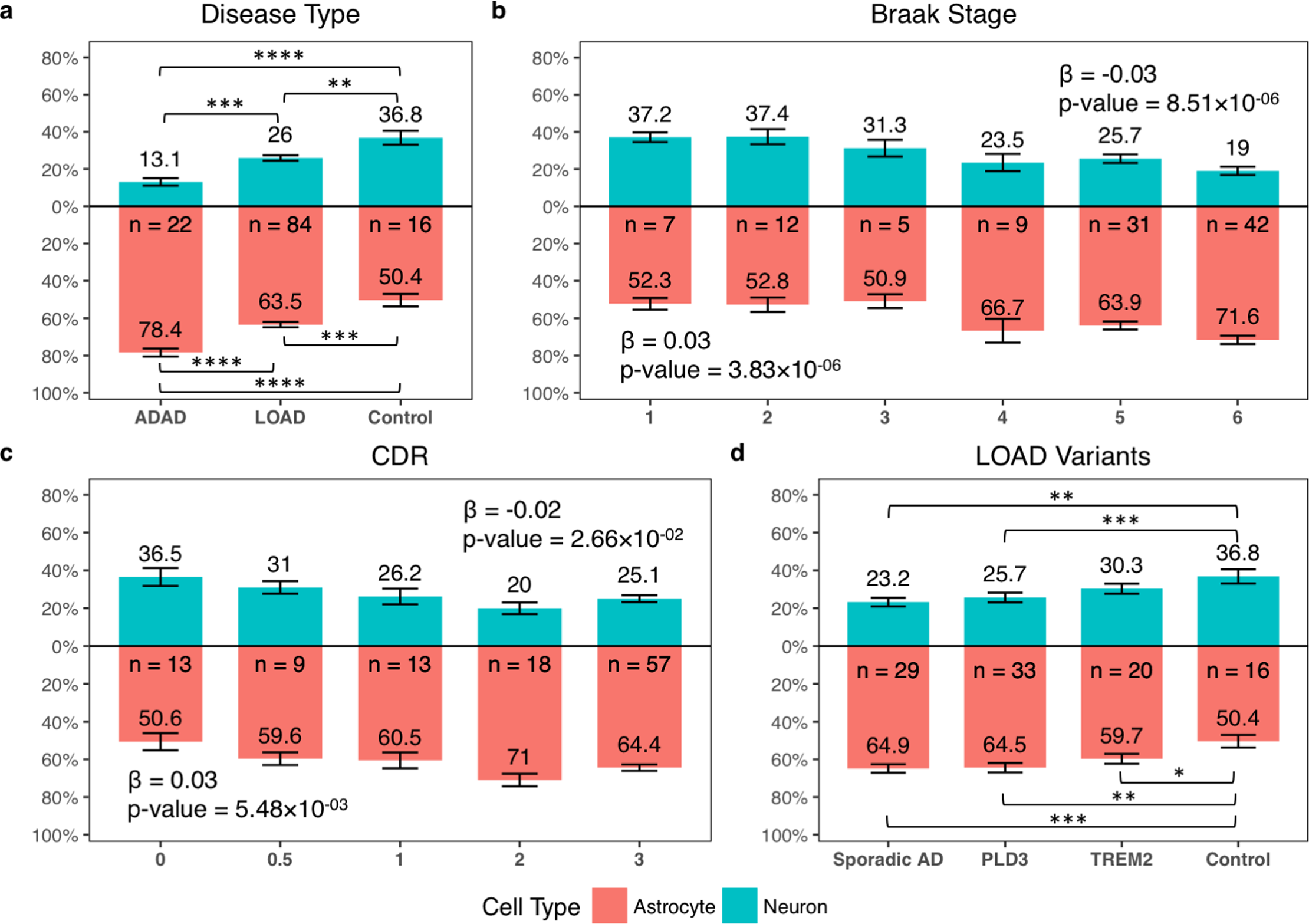
Neuron and astrocyte distributions from the DIAN and Knight-ADRC brains. **a)** Mean neuronal (blue) and astrocytic (red) proportions for carriers of pathogenic mutations in APP, PSEN1 or PSEN2 (ADAD), late-onset AD (LOAD) and neuropath-free controls (bars indicate standard deviations). Neuronal and astrocytic proportions plotted against **b)** Braak Staging; **c)** by Clinical Dementia Rating. **d)** Cell-type distributions for carriers of AD genetic risk factors. Lines indicate significance levels (*P< 0.05; ** P< 0.01; *** P< 1.0×10^−3^; **** P< 1.0×10^−4^).

**Table 3.** Cellular population structure altered in the parietal lobe from AD brains in the DIAN study and Knight-ADRC brain bank.

| **Disease Status** | **Sample Size** | **Neuron** |  | **Astrocyte** |  | **Oligodendrocyte** |  | **Microglia** |  |
| --- | --- | --- | --- | --- | --- | --- | --- | --- | --- |
| **AD Status** | N | Effect | P-value | Effect | P-value | Effect | P-value | Effect | P-value |
| AD<sup>a</sup> vs Control | 122 | -0.11 | $5.52 \times 10^{-04}$ | 0.14 | $2.48 \times 10^{-05}$ | -0.03 | $6.5 \times 10^{-02}$ | $-2.64 \times 10^{-03}$ | $2.49 \times 10^{-01}$ |
| ADAD vs Control | 38 | -0.19 | $3.94 \times 10^{-07}$ | 0.24 | $1.57 \times 10^{-10}$ | -0.04 | $8.5 \times 10^{-03}$ | -0.01 | $7.77 \times 10^{-05}$ |
| LOAD vs Control | 100 | -0.09 | $5.67 \times 10^{-03}$ | 0.12 | $3.34 \times 10^{-04}$ | -0.02 | $1.06 \times 10^{-01}$ | $-1.70 \times 10^{-03}$ | $4.57 \times 10^{-01}$ |
| ADAD vs LOAD |  |  |  |  |  |  |  |  |  |
| Braak matched | 42 | -0.08 | $1.03 \times 10^{-02}$ | 0.11 | $9.26 \times 10^{-04}$ | -0.03 | $7.1 \times 10^{-02}$ | $-1.46 \times 10^{-03}$ | $7.01 \times 10^{-01}$ |
| Braak corrected | 91 | -0.09 | $4.71 \times 10^{-03}$ | 0.11 | $5.24 \times 10^{-04}$ | -0.02 | $1.77 \times 10^{-01}$ | $-2.41 \times 10^{-03}$ | $4.25 \times 10^{-01}$ |
| CDR corrected | 94 | -0.12 | $2.11 \times 10^{-03}$ | 0.13 | $6.29 \times 10^{-04}$ | -0.02 | $3.8 \times 10^{-01}$ | $-3.11 \times 10^{-03}$ | $2.41 \times 10^{-01}$ |
| **Clinical Dementia Rating** |  |  |  |  |  |  |  |  |  |
| AD<sup>a</sup> and Controls | 110 | -0.02 | $2.66 \times 10^{-02}$ | 0.03 | $5.48 \times 10^{-03}$ | -0.01 | $2 \times 10^{-01}$ | $-4.63 \times 10^{-04}$ | $4.77 \times 10^{-01}$ |
| ADAD and Controls | 26 | -0.08 | $4.12 \times 10^{-04}$ | 0.11 | $1.78 \times 10^{-07}$ | 0.01 | $4.03 \times 10^{-03}$ | $-1.55 \times 10^{-03}$ | $1.75 \times 10^{-08}$ |
| LOAD and Controls | 100 | -0.02 | $3.22 \times 10^{-02}$ | 0.03 | $7.01 \times 10^{-03}$ | -0.01 | $1.81 \times 10^{-01}$ | $-4.64 \times 10^{-04}$ | $5.11 \times 10^{-01}$ |
| **Braak Staging** |  |  |  |  |  |  |  |  |  |
| AD<sup>a</sup> and Controls | 106 | -0.03 | $8.51 \times 10^{-06}$ | 0.03 | $3.83 \times 10^{-06}$ | $-4.24 \times 10^{-03}$ | $2.04 \times 10^{-01}$ | $-2.52 \times 10^{-04}$ | $6.81 \times 10^{-01}$ |
| ADAD and Controls | 33 | -0.05 | $2.37 \times 10^{-05}$ | 0.06 | $2.45 \times 10^{-05}$ | -0.01 | $2.29 \times 10^{-01}$ | $-7.2 \times 10^{-04}$ | $4.89 \times 10^{-01}$ |
| LOAD and Controls | 88 | -0.03 | $7.41 \times 10^{-04}$ | 0.03 | $4.63 \times 10^{-04}$ | $-3.72 \times 10^{-03}$ | $3.29 \times 10^{-01}$ | $-1.66 \times 10^{-04}$ | $7.86 \times 10^{-01}$ |

<sup>a</sup> AD includes both autosomal dominant AD (ADAD) and late-onset AD (LOAD).
The cellular population structure was inferred using the ssNMF method. Effects and p-values for the association with disease status, clinical dementia rating and Braak staging using generalized mixed models. We identified similar trends with approximately the same significance levels.

Next, we compared the cell proportion of LOAD vs ADAD and found that the cell composition differs between them. We firstly selected the LOAD brains (N = 25) to match the Braak staging distribution of ADAD brains (N = 17). The ADAD brains showed a significant decreased neuronal proportion compared to LOAD brains (β = −0.08; p = 1.03×10^−02^; Table 3), and increased astrocytes (β = 0.11; p = 9.26×10^−04^; Table 3). Then, we analyzed the entire Knight-ADRC LOAD brains, by extending the model to correct for Braak stages. We also observed significant decreased neurons (β = −0.09; p = 4.71×10^−03^; Table 3; Fig 3a; Table S9) and increased astrocytes (β = 0.11; p = 5.24×10^−04^; Table 3; Fig 3a; Table S9 in ADAD brains compared to LOAD. We observed the same cellular differences when we corrected for CDR at death (β = −0.12; p = 2.11×10^−03^ for neurons and β = 0.13; p = 6.29×10^−04^ for astrocytes; Table 3; Fig 3bc). In summary, our results indicate that ADAD individuals present a higher neuronal death even in the same stage of the disease, suggesting that in ADAD neuronal death play a more important role in pathogenesis than sporadic AD, in which other factors such as inflammation or immune response may be involved.

### Specific genetic variants confer a distinctive cell composition profile

A variety of genetic variants increase risk of LOAD; however, it is unclear if the cellular mechanisms are the same across these distinct risk factors. Therefore, we tested the hypothesis that distinct genetic causes of LOAD have characteristic cellular population signatures.

We initially ascertained the effect of *APOE* ε4 on the cell-type composition. We observed a significant decrease in neurons (β = −0.06 for each of the ε4 alleles; p = 9.91×10^−03^) and increase of astrocytes (β = 0.10; p = 4.15×10^−02^) from the TC included in the Mayo Clinic dataset (**Table S10; Fig 4a; Fig S9a**). This finding was replicated when we performed a multi-area analysis of the MSBB dataset (β = −0.03; p = 2.75×10^−04^ and β = 0.04; p = 8.06×10^−06^ for neurons and astrocytes respectively; Table 4; Fig 4a; **Table S10; Fig S9a**). Given the strong risk conferred by the *APOE* ε4 allele [19], we studied its effects on the cell-type composition by restricting our analysis to AD brains. We observed a significant association in the multi-area analysis of the MSBB dataset, with the same effect size for the neurons as the observed when we analyzed both affected and control brains (p = 1.60×10^−02^; Table 4; Fig 4b; **Table S11; Fig S9b**) and also a significant increase in astrocytes (β = 0.03; p = 1.03×10^−02^; Table 4; Fig 4b; **Table S11; Fig S9b**). We also observed a significant decrease in neurons proportion (β = −0.06; p = 2.11×10^−02^; Table 4; Fig 4c) when we analyzed the LOAD and control brains from the Knight-ADRC. When we restricted the analysis to AD brains from the Knight-ADRC and compared the *APOE* ε4 carriers (N = 46) to non-carriers (N = 41) we also observed decreased neurons (β = −0.06; p = 2.69×10^−02^; Table 4; Fig 4d).

**Fig 4.**
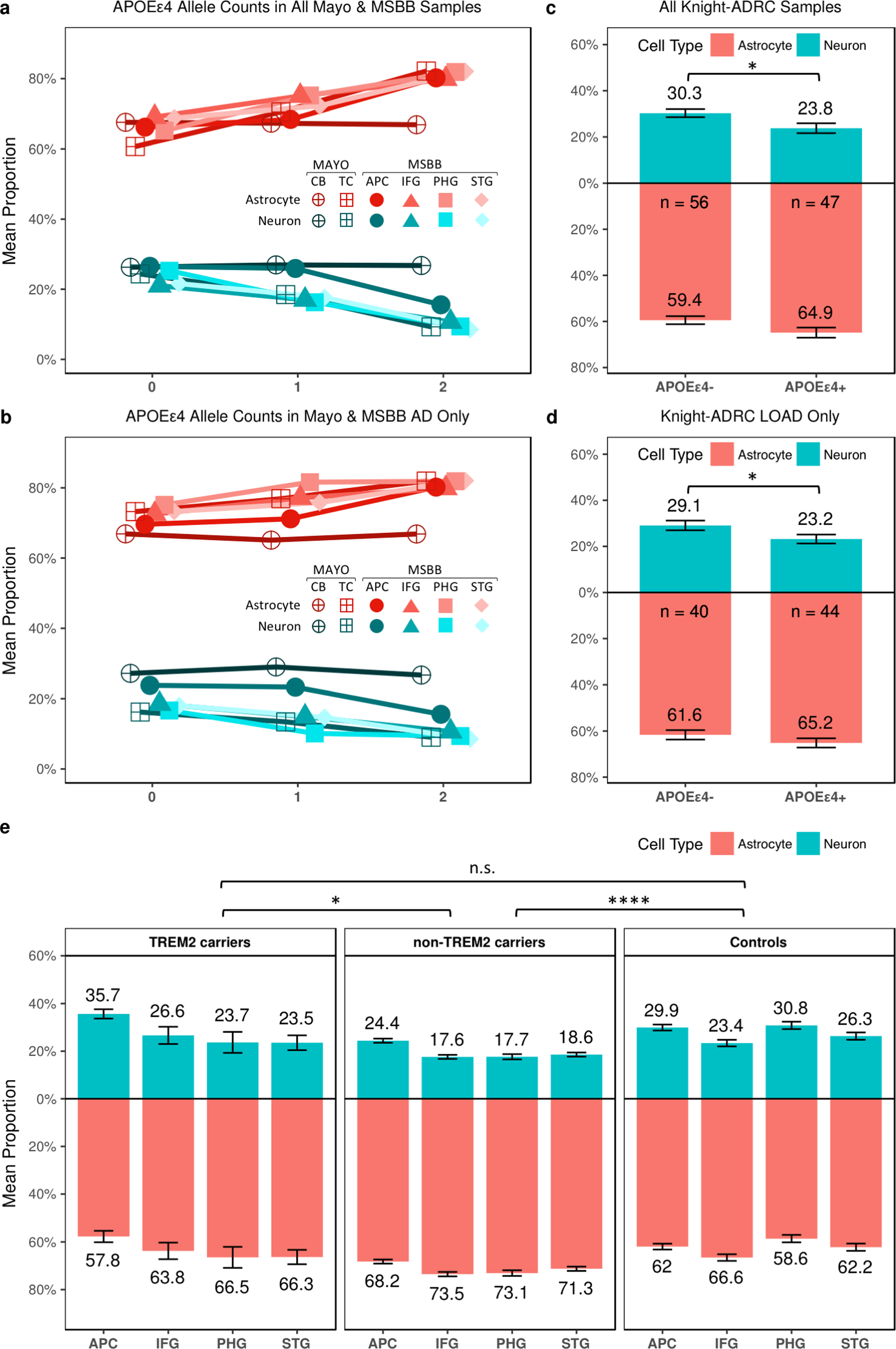
Effect of the APOE ε4 allele and TREM2 coding variants on the cellular population structure. Mean neuronal (blue) and astrocytic (red) proportions for **a)** AD cases and controls in the Knight-ADRC brains categorized by *APOE* ε4 carriers vs. non-carriers and **b)** AD cases of Knight-ADRC brain bank (bars indicate standard deviations). **c)** AD cases and controls in the Mayo Clinic and MSBB **d)** AD cases in the Mayo Clinic and MSBB. **e)** Neuronal (blue) and astrocyte (red) distributions for samples included in the Mount Sinai brain bank stratified by *TREM2* genetic status. APC: Anterior Prefrontal Cortex; STG: Superior Temporal Gyrus; PHG: Parahippocampal Gyrus; IFG: Inferior Frontal Gyrus; (n.s. P > 0.05; * P< 0.05; **** P< 1.0×10^−4^).

**Table 4.** Gene specific cellular proportion analysis for Knight-ADRC and Mount Sinai Brain Bank studies

| Variant Carriers | Sample Size | Neuron |  | Astrocyte |  | Oligodendrocyte |  | Microglia |  |
| --- | --- | --- | --- | --- | --- | --- | --- | --- | --- |
|  | N | Effect | P-value | Effect | P-value | Effect | P-value | Effect | P-value |
| **Knight-ADRC** |  |  |  |  |  |  |  |  |  |
| *PLD3* vs Control | 49 | -0.1 | $1.6 \times 10^{-04}$ | 0.13 | $2.84 \times 10^{-03}$ | -0.03 | $6.17 \times 10^{-02}$ | $7.05 \times 10^{-04}$ | $7.89 \times 10^{-01}$ |
| *TREM2* vs Control | 36 | -0.07 | $7.93 \times 10^{-02}$ | 0.11 | $1.05 \times 10^{-02}$ | -0.03 | $4.9 \times 10^{-02}$ | $1.65 \times 10^{-03}$ | $5.84 \times 10^{-01}$ |
| Sporadic AD vs Control | 45 | -0.11 | $5.45 \times 10^{-03}$ | 0.13 | $2.95 \times 10^{-04}$ | -0.02 | $4.55 \times 10^{-01}$ | $-3.48 \times 10^{-03}$ | $1.13 \times 10^{-01}$ |
| *APOEε4+* vs *APOEε4-* LOAD cases and controls | 100 | -0.06 | $2.11 \times 10^{-02}$ | 0.05 | $5.35 \times 10^{-02}$ | 0.01 | $3.72 \times 10^{-01}$ | $-8.09 \times 10^{-04}$ | $6.31 \times 10^{-01}$ |
| *APOEε4+* vs *APOEε4-* LOAD cases only | 84 | -0.06 | $2.69 \times 10^{-02}$ | 0.03 | $2 \times 10^{-01}$ | 0.03 | $1.4 \times 10^{-02}$ | $-8.31 \times 10^{-04}$ | $6.21 \times 10^{-01}$ |
| **Mount Sinai Brain Bank - Multi-region** |  |  |  |  |  |  |  |  |  |
| AD *TREM2* carriers vs Control | 301 | -0.03 | $3.57 \times 10^{-01}$ | 0.03 | $3.19 \times 10^{-01}$ | $-2.08 \times 10^{-03}$ | $7.87 \times 10^{-01}$ | $-2.68 \times 10^{-03}$ | $8.67 \times 10^{-02}$ |
| AD non-carriers *TREM2* vs Control | 882 | -0.07 | $1.91 \times 10^{-08}$ | 0.08 | $1.25 \times 10^{-08}$ | $-3.36 \times 10^{-03}$ | $4.79 \times 10^{-01}$ | $-2.89 \times 10^{-04}$ | $7.97 \times 10^{-01}$ |
| AD *TREM2* vs AD non-*TREM2* | 673 | 0.05 | $1.98 \times 10^{-02}$ | -0.05 | $1.58 \times 10^{-02}$ | $2.12 \times 10^{-03}$ | $7.76 \times 10^{-01}$ | $-2.13 \times 10^{-03}$ | $1.74 \times 10^{-01}$ |
| CDR corrected | 673 | 0.04 | $5.83 \times 10^{-02}$ | -0.04 | $4.46 \times 10^{-02}$ | $1.68 \times 10^{-03}$ | $8.19 \times 10^{-01}$ | $-1.92 \times 10^{-03}$ | $2.22 \times 10^{-01}$ |
| Braak corrected | 642 | 0.05 | $1.3 \times 10^{-02}$ | -0.05 | $2.7 \times 10^{-02}$ | $-1.82 \times 10^{-03}$ | $8.13 \times 10^{-01}$ | $-2.66 \times 10^{-03}$ | $1.28 \times 10^{-01}$ |
| Mean plaque counts corrected | 673 | 0.05 | $2 \times 10^{-02}$ | -0.05 | $1.59 \times 10^{-02}$ | $1.73 \times 10^{-03}$ | $8.15 \times 10^{-01}$ | $-2.2 \times 10^{-03}$ | $1.5 \times 10^{-01}$ |
| *APOEε4* counts all samples | 556 | -0.03 | $2.75 \times 10^{-04}$ | 0.04 | $8.06 \times 10^{-06}$ | $-4.33 \times 10^{-04}$ | $4.31 \times 10^{-01}$ | -0.01 | $2.42 \times 10^{-03}$ |
| *APOEε4* counts AD cases | 225 | -0.03 | $1.60 \times 10^{-02}$ | 0.03 | $1.03 \times 10^{-02}$ | $-2.07 \times 10^{-04}$ | $8.20 \times 10^{-01}$ | $-3.46 \times 10^{-03}$ | $3.29 \times 10^{-01}$ |

Next, we analyzed the cellular composition in *PLD3* carriers (N = 33). *PLD3* carriers exhibited significantly decrease of neurons compared to controls (β = −0.10; p = 1.60×10^−04^; Fig 3d) and a significant increase in astrocytes (β = 0.13; p = 2.84×10^−03^; Table 4; Fig 3d). Sporadic AD non-carriers cases also exhibited significantly decrease of neurons compared to controls (β = −0.11; p = 5.45×10^−03^) and significant increase of astrocytes (β = 0.13; p = 2.95×10^−04^; Table 4; Fig 3d). The cell proportion between sporadic AD non-carriers and *PLD3* carriers did not show any significantly difference (p > 0.05).

Finally, we performed similar analyses with *TREM2* carriers. *TREM2* is involved in the immune response and its role in amyloid-β deposition or clearance remain controversial [67]. Our analysis on the Knight-ADRC data showed significantly increased astrocytes in AD affected *TREM2* carriers (N = 20) compared to controls (β = 0.11; p = 1.05×10^−02^; Table 4; Fig 3d). Despite *TREM2* carriers presented lower neuron proportion compared to controls, this difference was not statistically significant (p>0.05; Table 4; Fig 3d). We analyzed whether the *TREM2* carriers provided sufficient power to detect a significant association. Our empirical estimates showed that *TREM2* sample size provides 96% of power to detect an association with an effect size comparable to that observed for sporadic AD (β = −0.11). We also investigated whether the cellular proportion of the eleven *TREM2* carriers in the MSBB dataset. The multi-region analysis showed *TREM2* carriers do not show a significant difference in neurons compared to controls (p > 0.05; Table 4; Fig 4e), whereas in the AD *TREM2* non-carriers the neuronal and astrocytic proportions are significantly different from controls (β = −0.07; p = 1.91×10^−08^ and β = 0.08; p = 1.25×10^−08^ respectively; Table 4; Fig 4e).

In fact, our analyses indicate that *TREM2* carriers have a unique cellular brain composition distinct than the other AD cases. *TREM2* brains showed significantly higher neurons (β = 0.05; p = 1.98×10^−02^) and significantly decreased astrocytes than the AD *non-*carries (β = −0.05; p = 1.58×10^−02^; Table 4). The distribution of CDR, mean number of amyloid plaques and Braak staging do not differ between strata. Nonetheless, we verified that the cellular proportions were still significant after correcting for each of those variables (Table 4). These results suggested that the mechanism that lead to disease in *TREM2* carriers is less neuron-centric than in the general AD population.

### Discussion

We have developed, optimized and validated a digital deconvolution approach to infer cell composition from bulk brain gene expression that integrates publicly available cell-type specific expression data while addressing the heterogeneity of the phenotypic differences of samples and technical characteristics of transcriptome ascertainment. We acknowledge that the accuracy of this platform might be affected by the phenotypic diversity of the reference panel or the disease-induced dysregulation of genes it includes. However, the deconvolution approach proved to be robust to the genes included in the reference panel, as we demonstrated that the proportions it inferred are not driven by the expression of any single gene. This platform produced reliable cell proportion estimates, as was shown by the evaluation of independent datasets of iPSC-derived neurons and microglia, mice cortical neurons (**Fig S4**) and simulated chimeric libraries.

We used this approach to deconvolve studies that include large number of neuropathologically defined AD and control brains with their transcriptome ascertained in distinct brain regions, and observed consistently significant neuronal loss and astrocytosis in the cerebral cortex. Compatible with other studies, we also identified that the altered cellular proportion is also significantly associated with decline in cognition and Braak staging [60]. In contrast, we did not identify a significant difference in the cellular population structure in the cerebellum, a region not affected in AD (Table 2; Fig 2a).

We generated RNA-seq data from brains carrying pathogenic mutations in *APP*, *PSEN1*, *PSEN2*, which cause alterations in Aβ processing and lead to ADAD, and also generated RNA-seq from brains of LOAD and neuropath-free controls. We observed altered cell composition in both ADAD and LOAD compared to controls. However, we identified that ADAD brains have a different cell-type composition than disease-stage-matched LOAD, as the ADAD has a significantly lower neuronal proportion and more pronounced astrocytosis. Based on our results, we would hypothesize that this change in Aβ processing of ADAD would leads to more direct to neuronal death than the pathological processes of LOAD. Similarly, decreased neurons and increased astrocytes were significantly associated with *APOEε4* allele. It has been reported APOE ε4 allele increase the risk for AD by affecting APP metabolism or Aβ clearance [15, 38], suggesting a direct link between APP metabolism and neuronal death.

In contrast, the analysis of the Knight-ADRC brains showed that the neuronal loss is less pronounced in *TREM2* carriers than in other LOAD cases. We replicated this finding in a multi-area analysis from the MSBB dataset. These results may implicate that *TREM2* risk variants lead to a cascade of pathological events that differ from those occurring in sporadic AD cases, which is also consistent with the known biology of *TREM2*. *TREM2* is involved in AD pathology through microglia mediated pathways, implicated on altered immune response and inflammation [18]. Recent studies in *TREM2* knock-out animals showed that fewer microglia cells were found surrounding Aβ plaques with impaired microgliosis [70]. Furthermore, *TREM2* deficiency was reported to attenuate tauopathy against brain atrophy [42]. We found no significant difference in the proportion of microglia between AD cases and controls. However, we found significantly decreased microglia in brains exhibiting pathological aging (**Table S7; Fig S6**), proving that these studies are sufficiently powered to identify significant differences. In any case, we cannot rule out the possibility of a change in the activation stage of microglia in these individuals. Overall, these results suggest that *TREM2* affects AD risk through a slightly different mechanism to that of ADAD or LOAD in general. Therefore, other pathogenic mechanisms should contribute to disease. We believe that a detailed modeling of immune response cells, reflecting the alternative microglia activation states, will generate more accurate profiles to elucidate the immune cell distribution in AD.

There is a large interest in the scientific community to use brain expression studies to try to identity novel pathogenic mechanism in AD and to identify novel therapeutic targets. These efforts are generating a large amount of bulk RNA-seq data, as single-cell RNA (scRNA-seq) from human brain tissue in large sample size is not feasible. Single-cell sorting needs to be performed with fresh tissue [33], which restrains the analysis of highly characterized fresh-frozen brains collected by AD research centers. Our results indicate that digital deconvolution methods can accurately infer relative cell distributions from brain bulk RNA-seq data. Having this approach validated for AD can have an important impact in the community, because digital deconvolution analyses 1) can reveal distinct cellular composition patterns underlying different disease etiologies 2) can provide additional insights about the overall pathologic mechanisms underlying different mutations carriers for variants as in genes such as *TREM2*, *APOE, APP, PSEN1* and *PSEN2*) can correct the effect that altered cell composition and genetic statuses have in addition to downstream transcriptomic analyses and lead to novel and informative results. 4) can help the analysis of highly informative frozen brains collected over the years.

In conclusion, our study provides a reliable approach to enhance our understanding of the fundamental cellular mechanisms involved in AD and enable the analysis of large bulk RNA-seq data that may lead to novel discoveries and insights into neurodegeneration.

## Acknowledgements

We thank all participants and their families for their commitment and dedication to helping advance research into the early detection and causation of AD; and the Knight-ADRC and DIAN research and support staff at each of the participating sites for their contributions to this study.

This work was supported by grants from the National Institutes of Health (R01-AG044546, P01-AG003991, RF1AG053303, R01-AG035083, and R01-NS085419), and the Alzheimer’s Association (NIRG-11-200110). This research was conducted while CC was a recipient of a New Investigator Award in Alzheimer’s disease from the American Federation for Aging Research. CC is a recipient of a BrightFocus Foundation Alzheimer’s Disease Research Grant (A2013359S). This work was supported in part by NIH K01AG046374 awarded to CMK, and also the Tau Consortium (CMK). JDD is supported by the Brain and Behavior Research Foundation and the NIH (R01NS102272). The recruitment and clinical characterization of research participants at Washington University were supported by NIH P50 AG05681, P01 AG03991, and P01 AG026276.

Data collection and sharing for this project was supported by The Dominantly Inherited Alzheimer’s Network (DIAN, U19AG032438) funded by the National Institute on Aging (NIA), the German Center for Neurodegenerative Diseases (DZNE), Raul Carrea Institute for Neurological Research (FLENI), Partial support by the Research and Development Grants for Dementia from Japan Agency for Medical Research and Development, AMED, and the Korea Health Technology R&D Project through the Korea Health Industry Development Institute (KHIDI). This manuscript has been reviewed by DIAN Study investigators for scientific content and consistency of data interpretation with previous DIAN Study publications. We acknowledge the altruism of the participants and their families and contributions of the DIAN research and support staff at each of the participating sites for their contributions to this study.

We would like to thank the operations staff at the Elizabeth H. and James S. McDonnell III Genome Institute at Washington University with their assistance in constructing the RNA-seq libraries and generating sequence data for our project. This work was also supported by accessing to equipment made possible by the Hope Center for Neurological Disorders and the Departments of Neurology and Psychiatry at Washington University School of Medicine. We also thank Allison M. Lake for her comments and suggestions.

The results published here are in whole or in part based on data obtained from the AMP-AD Knowledge Portal accessed at doi:10.7303/syn2580853. Study data were provided by the following sources: The Mayo Clinic Alzheimer’s Disease Genetic Studies, led by Dr. Nilufer Taner and Dr. Steven G. Younkin, Mayo Clinic, Jacksonville, FL using samples from the Mayo Clinic Study of Aging, the Mayo Clinic Alzheimer’s Disease Research Center, and the Mayo Clinic Brain Bank. Data collection was supported through funding by NIA grants P50 AG016574, R01 AG032990, U01 AG046139, R01 AG018023, U01 AG006576, U01 AG006786, R01 AG025711, R01 AG017216, R01 AG003949, NINDS grant R01 NS080820, CurePSP Foundation, and support from Mayo Foundation. Study data includes samples collected through the Sun Health Research Institute Brain and Body Donation Program of Sun City, Arizona. The Brain and Body Donation Program is supported by the National Institute of Neurological Disorders and Stroke (U24 NS072026 National Brain and Tissue Resource for Parkinson’s Disease and Related Disorders), the National Institute on Aging (P30 AG19610 Arizona Alzheimer’s Disease Core Center), the Arizona Department of Health Services (contract 211002, Arizona Alzheimer’s Research Center), the Arizona Biomedical Research Commission (contracts 4001, 0011, 05-901 and 1001 to the Arizona Parkinson’s Disease Consortium) and the Michael J. Fox Foundation for Parkinson’s Research. These data were generated from postmortem brain tissue collected through the Mount Sinai VA Medical Center Brain Bank and were provided by Dr. Eric Schadt from Mount Sinai School of Medicine. These data were generated by Kristen Brennand, a New York Stem Cell Foundation - Robertson Investigator. This work was supported by the Brain and Behavior Research Foundation, NIH grant R01 MH101454 and the New York Stem Cell Foundation. We analyzed iPSC-derived microglia RNA-seq data funded by NIH U01AG046170.

